# Bias in data-driven estimates of the replicability of univariate brain-wide association studies

**DOI:** 10.1101/2023.09.21.558661

**Authors:** Charles D. G. Burns, Alessio Fracasso, Guillaume A. Rousselet

## Abstract

Recent studies have used big neuroimaging datasets to answer an important question: how many subjects are required for reproducible brain-wide association studies? These data-driven approaches could be considered a framework for testing the reproducibility of several neuroimaging models and measures. Here we test part of this framework, namely estimates of statistical errors of univariate brain-behaviour associations obtained from resampling large datasets with replacement. We demonstrate that reported estimates of statistical errors are largely a consequence of bias introduced by random effects when sampling with replacement close to the full sample size. We show that future meta-analyses can largely avoid these biases by only resampling up to 10% of the full sample size. We discuss implications that reproducing mass-univariate association studies requires tens-of-thousands of participants, urging researchers to adopt other methodological approaches.

## Introduction

The question of scientific reliability of brain-wide association studies (BWAS) was brought to the attention of many^1,2^ by Marek, Tervo-Clemmens *et al.*^3^, reigniting discussions^4–7^ about the ongoing reproducibility crisis in neuroscience and psychology^8–12^. Reproducibility is often used as an umbrella term covering replication, namely obtaining similar results by applying the same methods to new data^13,14^. Independent researchers are failing to replicate BWAS^15,16^, suggesting that such findings are untrustworthy. Our trust in the scientific field therefore relies on how well we can estimate the replicability of its findings.

For any given study design, recruiting more subjects is a reliable way of increasing the likelihood of replication by reducing sampling variability and increasing statistical power^8^. For a BWAS, aimed at characterising associations between brain measures and behaviours, collecting data is expensive. So, how many subjects are required? How do we know? Thousands are required^3,17^, according to data-driven approaches which quantify the issue of replicability for BWAS using large neuroimaging datasets from the Human Connectome Project^18^ (HCP with n = 1,200), the Adolescent Brain Cognitive Development study^19^ (ABCD with n = 11,874), and the UK Biobank^20^ (UKB with n = 35,735). Among numerous analyses in their study, Marek, Tervo-Clemmens *et al*.^3^ estimated statistical errors of univariate BWAS as a function of sample size. Such mass univariate BWAS often involve tens of thousands of correlations between a brain measure and a behavioural measure, most of which fail to replicate even with thousands of participants due to small underlying effect sizes. These replication failures can be explained by statistical errors of a study design such as false positive rates^11^ and low statistical power^8,21–25^. To estimate statistical errors in univariate BWAS, Marek, Tervo-Clemmens et al.^3^ treated a large sample as a population and then drew replication samples by resampling with replacement (henceforth resampling) from that population (Fig. 1). This avoids the expense of collecting new data by instead resampling from a large sample, while using effect sizes from the large sample as replication targets. Statistical power, for example, was then estimated as the proportion of significant effects in the full sample which were significant again in a resample, averaged over 1000 iterations for each resample size and significance threshold.

**Figure 1:**
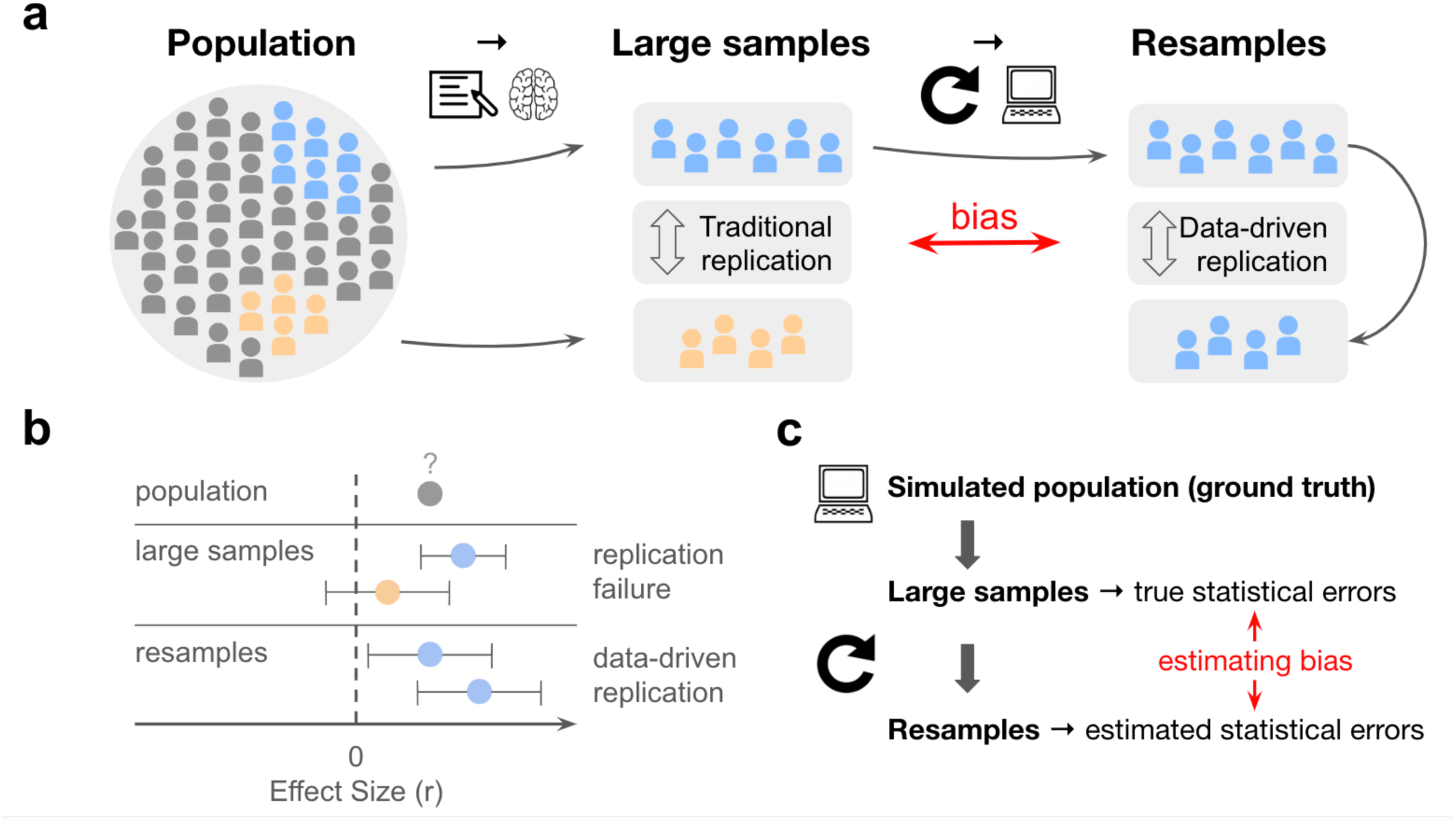
Overview of data-driven estimates of replicability. **a,** Schematic of large samples and resamples involved in Marek, Tervo-Clemmens et al.’s methods. **b,** Toy example of a single effect, where the population effect size (ρ), is unknown and estimated by large samples. Resamples and data-driven replications are used to estimate statistical errors underlying replication failures. Note that here replication failures are counted as a significant effect in a replication target (blue) being non-significant in a replication sample (orange), but better measures should be used in future work^26^. Confidence intervals illustrate uncertainty in effect size due to sampling variability and measurement reliability, showing larger intervals for smaller sample sizes (orange vs blue). **c**, Schematic of the current paper.

However, this data-driven method of estimating statistical errors might not generalise to the real-world scenario of repeated sampling from a population. Importantly, traditional replication involves comparing two large samples from a population, while data-driven replication involves comparing a large sample to its own resamples (Fig. 1 a). Data-driven simulations implicitly treat resampling from a large dataset as equivalent to sampling from a population, but these are not the same. As a result, we don’t know the difference between error estimates obtained from resamples and error estimates obtained from samples of a population – this difference we refer to as bias in statistical error estimates. Such bias may also be affecting other recent studies that relied on data-driven resampling methods to estimate statistical power^23,27,28^ and replicability^29^ in task-based fMRI methods, not just structural or functional associations. We therefore simulated data as ground truth to quantify bias in estimates of statistical errors.

## Results

### Resampling methods strongly bias statistical error estimates when there are no true effects

First, we simulated a large null sample with *n =* 1,000 subjects, each with 1,225 brain connectivity measures (random Pearson correlations) and a single behavioural measure (normally distributed across participants). We correlated each brain connectivity measure with the behaviour across all subjects to obtain 1,225 brain-behaviour correlations. Since brain connectivity estimates and behavioural factors were simulated independently from each other, any resulting brain-behaviour correlations were entirely random. In other words, the population effect size was null (*ρ* = 0). Data dimensions were chosen to be computationally feasible for reproducibility, however we invite readers to adjust these and re-run analyses using the openly available code (analyses recoded in R with supporting packages^30–32^ for open-source accessibility https://github.com/charlesdgburns/rwr/). We then resampled our null-sample for 100 iterations across logarithmically spaced sample size bins (*n* = 25, to 1,000) and estimated statistical errors, following the methods described in Marek, Tervo-Clemmens *et al*.^1^. Surprisingly, we saw the same trends of statistical errors and replicability as those reported by Marek, Tervo-Clemmens *et al*.^1^ but with random data (see Fig. 2), noticing strongly biased statistical power estimates.

**Figure 2:**
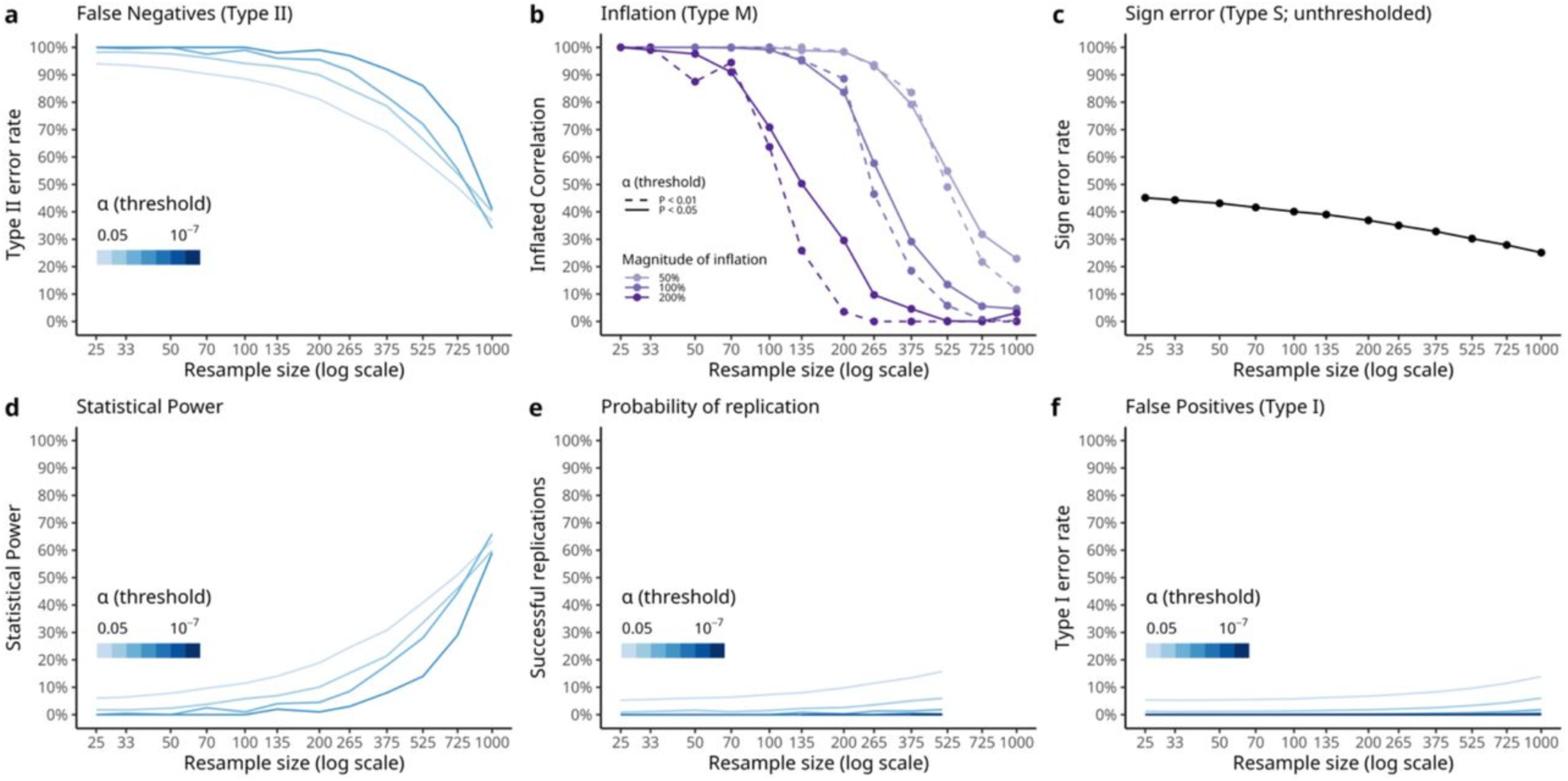
Estimated statistical errors and reproducibility of random noise (*ρ* = 0) We reproduce statistical error estimates after resampling from a simulated null-sample where observed significant effects are random (ρ = 0). These results are notably comparable to Fig. 3 in Marek et al.^3^ **a**, False negative rates computed relative to full sample size decrease as the resample size increases, across a range of significance thresholds which were passed after sampling 1,225 random correlations. **b**, For a given magnitude of inflation, when inflation rates are computed relative to effect sizes at the full sample size they fall to 0% as the resample size approaches the full sample size. **c**, Sign errors computed relative to signs of effects at the full sample size show a downwards trend as resample sizes increase from around chance level of 50% to 30%. **d**, Statistical power when computed relative to full sample size shows a strong upwards trend as resample sizes increase, reaching around 60% for all significance thresholds which were crossed when simulating 1,225 random effects (α=0.05 to 0.0001). **e**, Probability of replication, computed by resampling both equally sized ‘in-sample’ and ‘out-of-sample’ subsamples from the full dataset, stays low for small significance thresholds, but reaches above 10% for α=0.05. **f,** False positive rates, computed by counting correlations which are significant only in the resample but not in the full sample size, also reach about 10% for α=0.05 in the full sample size. Note that the darkest lines are drawn after random effects pass thresholds as low as α=10^−7^ after resampling from the full sample, which is lower than Bonferroni correction 0.05/1225 = 4×10^−5^.

Figure 2 suggests that the reported trends in estimated statistical errors do not depend on absolute sample size, but on the resample size relative to the full sample size. To establish a ground truth against which to assess these estimates, we consider what statistical error estimates we should expect if replication samples were drawn from a population (of infinite size) rather than resampled from a large sample. By repeatedly generating new null-samples, rather than resampling from a single null-sample, we verified that these statistical error estimates are indeed biased under the null as the resample size approaches the full sample size (Fig. 3). We also demonstrate that this bias stems from the act of resampling from a large sample, rather than how estimations are computed when comparing different large samples. For example, uncorrected (α = .05) statistical power at the full sample size (n = 1000) was estimated to be 63% when resampling (Fig 2 d), rather than the expected 5% obtained when generating new null-samples (Fig 3 d). One concern is that power is the most inflated while also being the most relevant for failed replications^8,21,22^, which could potentially result in misleading meta-science.

**Figure 3:**
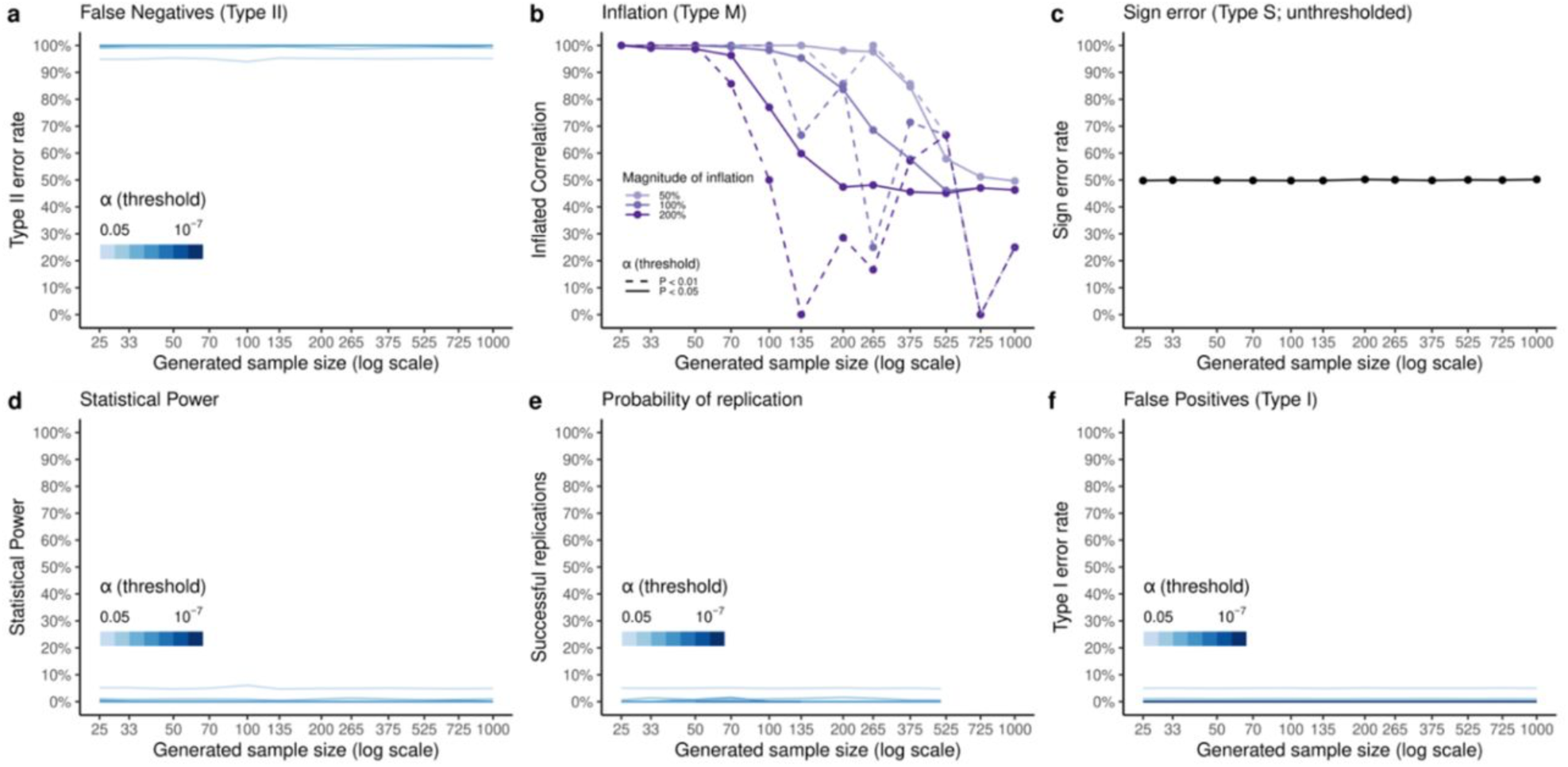
Expected estimates of statistical error and reproducibility of random noise (*ρ* = 0) We obtain estimates of statistical errors under the null by iteratively generating null-samples at increasing sample sizes (*n*=25,…,1,000) instead of resampling from a single null-sample, averaging estimates for each sample size over 100 simulations. This corresponds to sampling from an infinite-size population. **a,** estimated true false negative rates under the null are constant across sample sizes and equivalent to 1 - *α* for a given significance threshold (*α* = .05, .01, .001 plotted). **b,** since inflation rates are estimated as a proportion of replicated (same sign, and significant in both large sample and replication sample) correlations, we can expect these to be high for small sample sizes as the critical r for significance is higher, and so the likelihood of being inflated decreases as sample sizes increase. Since we are averaging across correlations, few of these will be very inflated while many will be less inflated so that on average this cancels out to 50% across inflation thresholds. **c,** we expect 50% sign errors regardless of sample size as the sign of a given correlation in a replication null-sample will be random. **d, e, f,** estimates of statistical power, probability of replication, and false positives are based on proportions of significant correlations in replication null-samples, so in each case the probability of a correlation being significant in a newly generated null-sample is exactly determined by the significance threshold (*α* = .05, …, 10^−7^).

### Compounding sampling variability underlies biased statistical errors under the null

To explain why bias arises under the null, we investigated the underlying brain-behaviour correlations used in the calculation of statistical errors. Here we focused on resampling at the full sample size (*n* = 1,000) where these biases are most dramatic. As indicated by the false positive rate (Fig. 2 f.), the null distribution of brain-behaviour correlations is not preserved when resampling at the full sample size (Fig. 4). Instead, resampling subjects and computing correlations again results in a distribution wider than expected (comparing Fig. 4 **a**. and **c**.). This is because resampling involves two sources of sampling variability, first at the level of the large sample and again for the resampled replication sample (Fig. 1 **a**). For instance, if a correlation in the large sample is randomly observed to be r = 0.11, then resampling participants and computing the same correlation again results in a correlation which varies around r = 0.11 (Fig. 4 **e**.).

**Figure 4:**
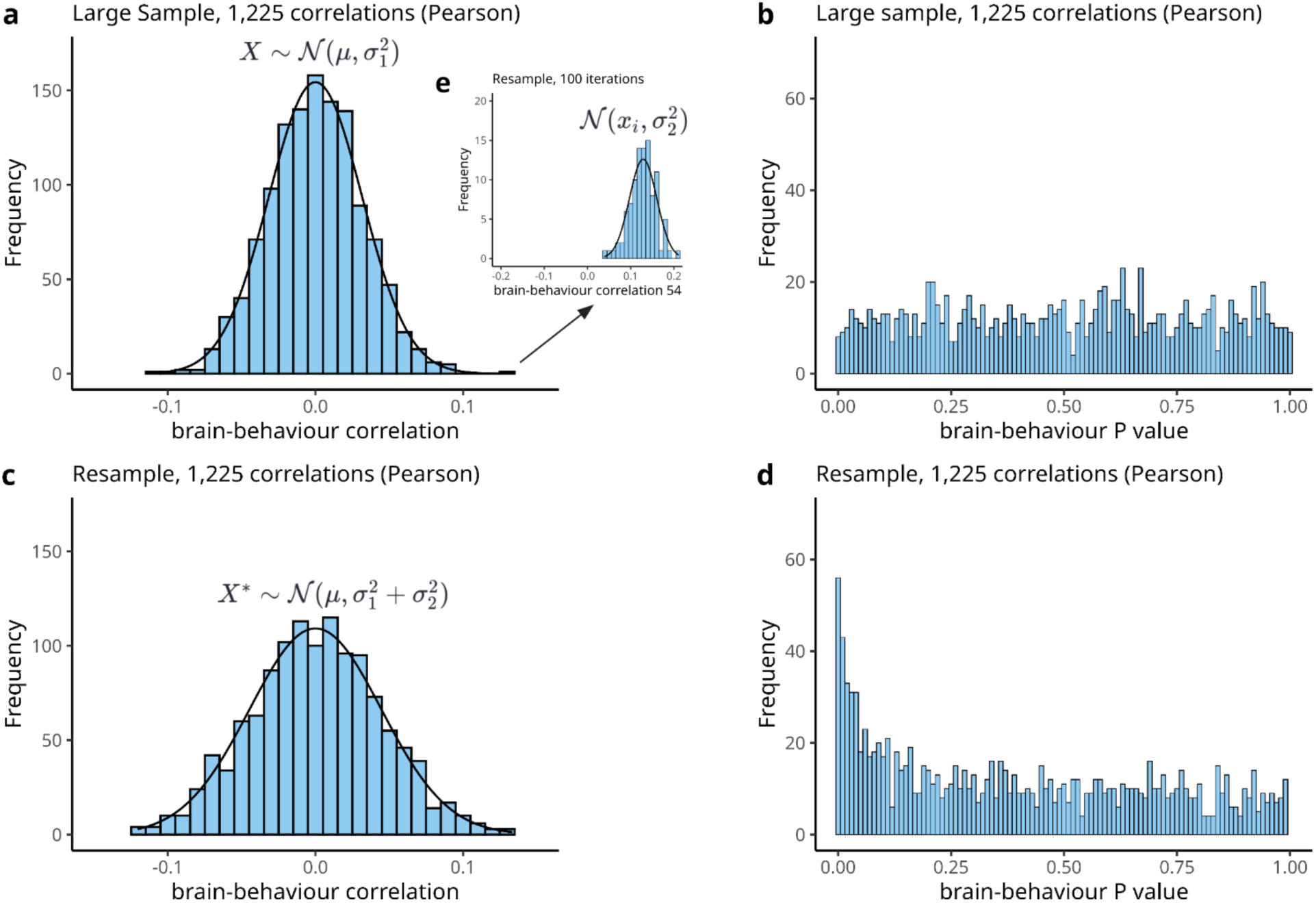
Null distributions under resampling with replacement (*ρ* = 0) **a**, Distribution of simulated random brain-behaviour correlations (1,225 total) treated as a large sample. There are n = 1,000 subjects for each random Pearson correlation, here compared to a Gaussian curve with mean µ = 0 and variance σ_1_^2^ = .001 drawn in black. **b**, We verify that the two-tailed *P* values of our 1,225 random brain-behaviour correlations are uniformly distributed. **c**, Distribution of all 1,225 brain-behaviour correlations computed after resampling subjects at full sample size (n = 1,000). This distribution is clearly wider than our large sample null distribution. The solid black line shows the distribution expected from an interaction of sampling variability (see **e**.), namely a Gaussian distribution with mean µ = 0 and variance σ_1_^2^ +σ_2_^2^ = .002. **d**, Distribution of two-tailed *P* values of all 1,225 brain-behaviour correlations computed after resampling at full sample size (n = 1,000). The distribution is inflated around 0 due to the wider tails in our null distribution. **e**, To help explain the widened distribution, we track the largest correlation observed in our original null-sample (r = 0.11), plotting the distribution of corresponding brain-behaviour correlations across the 100 iterations of resampling at the full sample size (n = 1,000). The solid black line represents a Gaussian with mean µ = 0.11 and variance σ_2_^2^ = .001. The interaction of variability across iterations (**e**) and variability in the large sample (**a**) results in the widened distribution (**c**) by additive variance.

We can formalise this mathematically as nested distributions^33^, or a convolution^34^ of two probability distributions, here approximating Pearson null distributions with normal distributions for analytical simplicity. It then follows that given a large sample *X ∼ N(µ, σ_1_*^2^), for each observation in our original sample, *x_i_* ∈ *X,* resampling participants and recomputing correlations corresponds to sampling from several distributions *X_i_* ∼ *N*(*x_i_*, *σ_2_*^2^), resulting in a final set of correlations distributed according to *X* ∼ N(µ, σ_1_^2^+ σ_2_*^2^). Note that *σ_1_*^2^ depends on the size of the large sample, while *σ_2_*^2^ is determined by the resample size.

The influence on statistical error estimates such as statistical power is two-fold. First, random correlations at the tail in a large sample are more likely to be at the tail of correlations in a resample (Fig, 4 **a** and **e**). This inflates power when estimated as the proportion of significant effects in the large sample which are significant again in the resample (1 – false negative rates). Second, increased sampling variability alone leads to a wider-than-expected distribution of correlations with more extreme tails. These more extreme tails lead to an inflation of *P* values close to 0 in our resample (compare Fig. 4 **b**. and **d**.) when calculated using a standard correlation function (e.g., ‘corr’ in MATLAB). We note that simply correcting for this widened null distribution will over-correct for bias in statistical error estimates when true effects are present (see Supplementary Information).

### Bias in ground truth simulations depends on statistical power of the full sample size

While we have shown clear biases when there are no true effects (ρ = 0), this does not imply that there will be biases when true effects are present (ρ ≠ 0). We therefore investigated bias in statistical error estimates when true effects are present, focusing on statistical power estimates. We note that Marek, Tervo-Clemmens *et al.*^3^ have already shown that the largest univariate effect is highly replicable even for moderate sample sizes, so there are at least some true BWAS effects in the real world. However, since the population effect size (ρ) remains unknown (Fig. 1. b.), we simulated a range of scenarios such that roughly 1% (an arbitrary but conservative proportion) of effect sizes were true effects. To have a clear ground truth separation between null and true effects when estimating error rate bias in each scenario, we therefore sampled 54,778 effects from a null (ρ = 0), and 500 effects from a true effect (ρ ≠ 0) for a total of 55,278 effects (333 choose 2; as in Marek, Tervo-Clemmens et al.,^3^) in each large sample (Fig. 5 a). We chose true effect sizes in each case so that they were evenly spaced from 0.1% to 99% statistical power at the full sample size (n = 1000; Fig. 5 b), as determined by an inverse power analysis (see Methods). Because the bias under the null is driven by the false rejection of null hypotheses, here we adopted a fixed significance threshold after Bonferroni correction, which controls for at least one false positive among all comparisons (family-wise error rate). We then estimated statistical power by resampling as in Marek, Tervo-Clemmens et al. (Fig 5 c.) and compared these to analytical power levels (Fig 5 b.) to quantify bias in statistical power estimates (Fig 5 d.).

**Figure 5:**
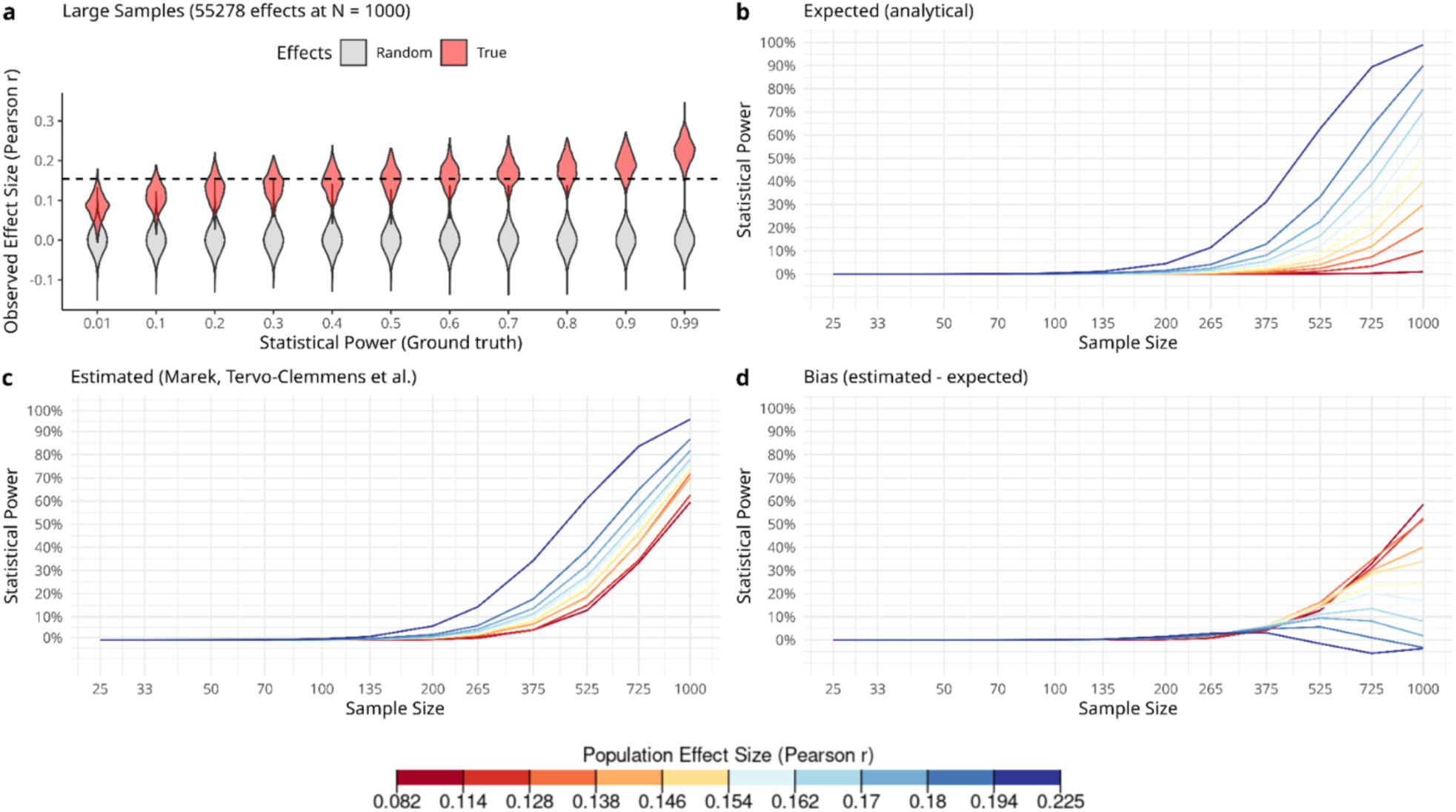
Bias in estimated statistical power depends on true statistical power. **a**, Simulated large samples are represented for each of the underlying power scenarios. For each scenario, grey violin plots show the distribution of 54,778 random effects and red violin plots represent the distribution of 500 effects sampled from an analytically derived true effect size (see Methods). The dashed line represents the critical Pearson *r* for a Bonferroni corrected significance level (α = 0.05/55278). **b**, For each scenario, we plot analytically derived power curves across sample size, representing the expected statistical power when repeatedly sampling from the population. Line colour represents each large sample scenario, noting the correspondence with power level at the full sample size (n = 1000). **c**, We estimated power across sample size by simulating the resampling methods in Marek, Tervo-Clemmens *et al.*^1^ using a Bonferroni corrected significance threshold. Line colour represents the ground truth statistical power of the study at full sample size (n = 1,000, α = 0.05/55278). **d**, To demonstrate bias across different sample sizes, we subtracted analytical power curves (panel b) from the estimated power (panel c), with lines coloured as in panel b. Note that underpowered large samples inflate statistical power estimates near the full sample size.

We show that the bias in statistical power estimates near the full sample size depend on the true statistical power of the large sample we resample from (Fig. 5). Very large samples are therefore required to accurately estimate statistical power for very small effects. Power estimates are inflated if the large sample is underpowered, but on the other hand a highly powered large sample may give conservative power estimates. This is driven by a subset of null and true effects in the large sample that are near the significance threshold; due to sampling variability, in a resample these small effects can easily cross the threshold and introduce type I and type II errors. Note that regardless of power at the full sample size, bias in statistical power is largely avoided when subsampling up to around 10% of the full sample size (see also Supplementary Information for subject-level simulations and larger effect sizes).

## Discussion

Accurately estimating reproducibility of scientific methods is critical for guiding researcher’s methodological decisions. Our results demonstrate that estimating statistical errors by resampling with replacement from random data results in large biases when the resample size approaches the full sample size. This is explained by compounding sampling variability of test statistics when resampling and its knock-on effects on estimated statistical errors. We further simulate data with true effects to show that statistical power is inflated when the true power of the large sample is low and slightly deflated when true power is high. This could lead to circular reasoning in cases where we must assume we have high statistical power before we can rely on the estimation that we have high statistical power. Lastly, we show that this bias is largely avoided when subsampling only up to 10% of the full sample size after Bonferroni correction. This 10% rule of thumb is consistent with the use of resampling techniques in a recent evaluation of statistical power and false discovery rates for genome-wide association studies with hundreds-of-thousands of participants^35^, as well as recommendations for 10-fold cross-validation to reduce upwards bias in prediction errors in machine-learning^36^.

What are the implications for the results presented by Marek, Tervo-Clemmens *et al*.^1^? Their estimates have been optimistic by an order of magnitude, implying that replicating mass univariate BWAS requires not thousands, but tens-of-thousands of participants. Revisiting their data, we can compare estimates at the full sample size, where we expect most bias, to estimates at 10% of the full sample size, where we expect no bias. For the strictly denoised Adolescent Brain Cognitive Development (ABCD) sample (*n* = 3,928), they report around 68% power after Bonferroni correction when resampling at the full sample size (Marek, Tervo-Clemmens *et al*.^1^ Fig. 4 **d**.). When subsampling from the UK Biobank with a full sample size of *n* = 32,572 Marek, Tervo-Clemmens *et al*.^1^ report around 1% power for *n* = 4,000 and *α* = 10*^−^*^7^. We therefore argue that the 68% power reported for the full ABCD sample (*n =* 3,928, *α* = 10*^−^*^7^) more likely reflects methodological bias, rather than increased signal after strict denoising of brain data. While the largest BWAS effects may be highly replicable with 4,000 participants, the average univariate BWAS effect is most likely not. Furthermore, our true effect simulations (Fig. 5) also indicate that the UK Biobank estimates at the full sample size (*n* = 32,572) could be more accurate, with an underlying power likely between 70% and 90% after Bonferroni correction. However, we note that our simulations and data-driven replication methods only account for sampling variability and do not account for measurement reliability^37^. Statistical power would be lower for less reliable measures, such as 5-minute resting-state functional connectivity compared to structural brain measures. Ultimately, traditional replication of mass univariate BWAS would require tens-of-thousands of individuals.

### Recommendations

We stress that our results only have direct implications for mass univariate association studies using the methods in Marek, Tervo-Clemmens *et al.*^3^. These methods are only a small subset among many options study associations between fMRI brain measures and behavioural measures, which warrants further investigations into replicability of studies using other methods. Here we therefore consider how some methodological choices could explain the lack of power in mass univariate BWAS and influence the replicability of neuroimaging studies^25^.

First, the study design can have a large influence on replicability by increasing statistical power. For example, inter-individual correlation studies offer “as little as 5%-10% of the power” of within-subject t-test studies with the same number of participants^4^, giving a power advantage to group-level designs^38^. Another way to increase power is to relieve the constraint on significance thresholds required to control for the many multiple comparisons involved in fMRI^39^. Studies can therefore focus on fewer pre-selected local brain regions^23,40^, or fewer measures which aggregate data across brain regions using networks^7,41–43^ or multivariate pattern analyses^3,44,45^. Studies which limit their analyses by pre-registration^9,46^ also tend to produce better powered studies^47^, suggesting that this practice encourages more careful study design.

Second, data processing in fMRI can have a large effect on reliability of brain measures and hence replicability^37^. For resting-state fMRI, two key confounds are head motion and global signal^48,49^ which should be carefully controlled for, noting that the choice of de-confounding methods can strongly influence resulting network measures^50,51^. A recent comparison of resting-state fMRI network analysis pipelines^48^ further showed that parcellation choice matters. Brain parcellation reduces the number of brain measures by grouping voxels into parcels if they share activity patterns in time. Pervaiz *et al.*^48^ recommend low-dimensional independent component analysis (ICA with D=50 regions) data-driven from the group of subjects within a dataset, in contrast to Marek, Tervo-Clemmens *et al.*’s^3^ choice of a pre-determined group-averaged brain parcellation^52^ (with D=333 regions). We note that both of these parcellations fail to account for individual level variations in resting state functional connectivity^53,54^. Future analyses could therefore benefit from recent methods^55^ which do account for variability of networks both between- and within-participants^53,56^. A key point here is that different processing choices make different underlying assumptions about brain data^57,58^, which can affect reliability. For example, different methods may be more or less sensitive to how much data is collected per participant compared to total data across participants. Having considered data processing pipelines which increase reliability, we note that researchers should also consider the reliability of behavioural measures^61^ when aiming for future studies with greater replicability.

Third, researchers should consider choosing prediction over explanation^62^, reporting results which are directly aimed at generalising to unseen data rather than relying on statistical inference in null-hypothesis significance testing (NHST) within a sample. Issues with NHST are well documented^26,63,64^ and combined with small sample sizes it leads to underpowered studies that report distorted effect sizes^8,11,22,65^. Another issue with NHST is that *P* values may be derived from inappropriate null models (as we saw in Fig. 4); choosing an appropriate null for brain-wide statistics^66^ is therefore yet another factor worth considering. These issues are largely avoided in a predictive framework, in which prediction accuracy of a model in a held-out-dataset provides a direct estimate of how well a model generalises. A key approach here is cross-validation, a machine learning strategy to prevent a model from overfitting on a single dataset. Recent studies have shown that when multivariate BWAS are cross-validated they report effect sizes that are replicable with only hundreds of participants^67,68^. Predictive models can therefore improve replicability by reporting effect sizes which are closer to true underlying effect sizes, however they should aim to do so while overcoming the challenge of interpretability^64^.

### Conclusive remarks

Our results show that previous data-driven estimates of statistical errors and replicability may have been optimistic. The implications are striking for univariate BWAS, but after considering the impacts of other methodological choices, it is also clear that investigations of replicability of wider BWAS methods are required. We urge such meta-analyses to evaluate their meta-analytic methods, for example with null data, so they may reliably evaluate the replicability of scientific methods used in research.

## Methods

### Simulating null data at subject level

We simulated random phenotype associations with simulated functional connectivity measures. We generated a null-sample with n = 1,000 subjects each with 1,225 edges (random Pearson correlations between 50 random time series) and a single behavioural factor (normally distributed across participants). We correlated each edge with the behaviour across all subjects to obtain 1,225 brain-behaviour correlations. By generating edge connectivity estimates and behavioural factors independently from each other, we ensured that any resulting brain-behaviour correlations are entirely random (ρ = 0), hence obtaining a sample where the null hypothesis is true (i.e., a null-sample).

### Estimating statistical errors

We closely followed the methods of Marek, Tervo-Clemmens *et al.*^3^, first running analyses on MATLAB using their code ‘abcd_edgewise_correlation_iterative_reliability_single_factor.m’ and ‘abcd_statisticalerrors.m’ (https://gitlab.com/DosenbachGreene/bwas). These analyses were then independently recoded in R with supporting packages^30–32^ for open-source accessibility https://github.com/charlesdgburns/rwr/. Notably, statistical error estimations involve two-tailed *P* values derived from parametric null distributions on a given resample size.

### Simulating ground truth data with known statistical power

At this stage we take a computationally more efficient approach and simulate summary statistics rather than subject-level data, which allows us to simulate many more true effect scenarios so we can compare estimates with true statistical errors. This approach also lets us increase the number of effects, so we now simulate samples with 55,278 (333 choose 2) effects, the number of resting-state functional connectivity measures which feature in Marek, Tervo-Clemmens *et al*. Fig 2^3^. The size of true effects was determined by an inverse power analysis with a fixed sample size (n = 1,000) and Bonferroni corrected significance threshold (α = 0.05/55278), using a Fisher z-transformation for calculating the critical Pearson r for a given power level (power = 1%, 10%, 20%, 30%, 40%, 50%, 60%, 70%, 80%, 90%, 99%). We derive the minimal effect size required to reach a given level of power, *r*_critical_, using the formula stated below, where *Z_α_* and *Z_β_* are the standard normal deviates for the significance threshold (α) and corresponding power level (1-β) respectively:

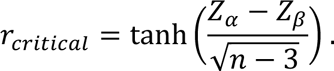

The Fisher z-transformation^70^, *F*(r) = atanh(r) = z was also used to sample Pearson correlations using the approximation that the z-statistic is asymptotically normally distributed with mean *F*(r) and standard deviation 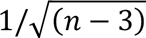. The z-statistic was then transformed into Pearson correlations to simulate brain-behaviour correlations. For each power level, we simulated a large sample by first drawing 55,278 random effects (ρ = 0) and afterwards replacing 500 of those with effects drawn from an infinite-sized population with true effect size ρ corresponding to the critical r for a given power level computed earlier. The choice of 500 effects being sampled from a true distribution was somewhat arbitrary, leading to a proportion true effects (∼1%) moderate enough to seem probable while sufficiently large to reduce noise in estimates of bias (see Methods section below). Note that while real world effect sizes of a single BWAS may vary, we instead simulate several effects with the same underlying effect size. This should not be an issue, since the statistical error summary statistics are given as an average across effects, so in this simulation we can think of the underlying effects having an average effect size according to a given power level. Similar estimates would be obtained if the underlying true effect sizes were varied but with an average effect size corresponding to each *r*_critical_.

### Simulating statistical power estimations from resampling with replacement

Given only summary statistics rather than individual subjects, we cannot resample participants and recompute *P* values, but instead also simulate obtaining estimates by resampling with replacement as in subject-level analyses^3^. First, we generate ground truth data with known statistical power, which we treat as a large sample. Then, we follow the implicit assumption that the observed effects in a large sample are the population effects: for a given resample size *n*, a resampled effect size was drawn from a normal distribution *N*(atanh(*r*\*), 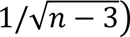, where r* is a given Pearson correlation from the large sample, and then Fisher z-transformed. We then derived *P* values from an uncorrected null distribution of Pearson correlations with degrees of freedom computed relative to the resample size. We resampled across the same range of sample sizes as in previous analyses (n = 25, … 1,000). We continued to estimate statistical power across 1,000 iterations of resampled brain-behaviour correlations as in Marek, Tervo-Clemmens et al.^1^, specifically as the proportion of significant effects in a large sample which were significant again in a resample (1 – false negative rates, with α = 0.05/55278). These were then compared to analytical power curves^69^ computed using Fisher z-transformations for varying sample sizes and effect sizes corresponding to critical r for power levels at the full sample size (n = 1,000) computed earlier.

## Data availability

No human data was collected for this study. R and MATLAB code used for data simulation, statistical analyses, and plotting is available on GitHub: https://github.com/charlesdgburns/rwr/.

## Ethics

Analyses described here were performed using randomly simulated data and were therefore not subject to ethical review.

## Acknowledgements

We thank editors and reviewers for their contributions to this manuscript. A.F. was supported by a grant from the Biotechnology and Biology research council (BBSRC, grant number: BB/S006605/1) and the Fundação Bial, Fundação Bial Grants Programme 2020/21, A-29315, number 203/2020, grant edition: G-15516.

## Authors’ contributions

C.D.G.B.: Conceptualisation, design, implementation, analysis, interpretation, writing - original draft. A.F.: Interpretation of results, writing - review & editing. G.A.R.: Conceptualisation, design, interpretation, writing - review & editing, supervision.

