## Supplementary Information for "Bias in data-driven estimates of the replicability of univariate brain-wide association studies"

**Can we adjust P values?**

Based on the compounding sampling variability and nested gaussians discussed in the main text, we consider calculating two-tailed *P* values using a parametric estimation of the resulting sampling distribution after resampling, approximated by $N(\mu, \sigma_1^2 + \sigma_2^2)$. The primary variance, $\sigma_1^2$, depends on the full sample size and secondary variance, $\sigma_2^2$, depends on the resample size, each being derived from the variance of the null distribution of Pearson correlations for the full sample size and resample size respectively. For a given sample size *n*, the null distribution of Pearson correlations, *r*, is described by the probability density function *P(r,n)*:

$$P(r, \mathrm{n}) = \frac{(1 - r^2)^{(n-4)/2}}{Beta(1/2, (n-2)/2)}$$

For each *n* we can then compute two-tailed *P* values for each correlation in our resample, r*, by:

$$P(X > |r^*|) = 2 \cdot \int_{|r*|}^{\infty} N(0, \sigma_1^2 + \sigma_2^2) dx$$

To visually check the adjusted *P* values and the validity of our gaussian approximation, we plot two sets of *P* value distributions, resampling from a full sample size of n = 1,000 at all the resample sizes, from n=25 up to the full sample size (Supplementary Figures 1 and 2). We do this separately for unadjusted *P* values, as these were calculated in Marek, Tervo-Clemmens *et al.*[1] and for the adjusted *P* values calculated here.

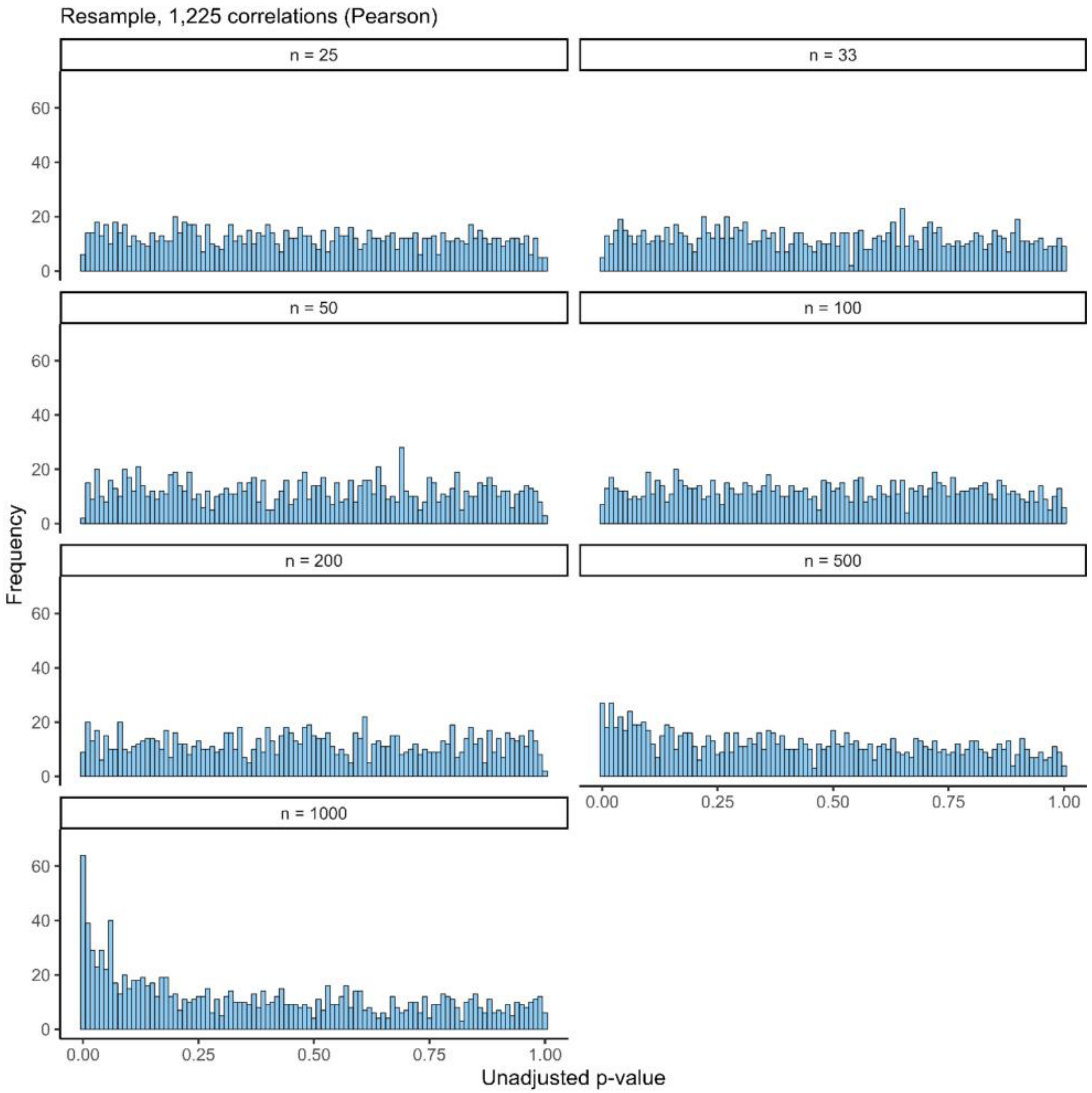


Supplementary Figure 1: **Distributions of unadjusted *P* values under different resample sizes.** Note that the full sample size is $n$ = 1,000, hence we resample from 2.5% of the full sample size to 100%. The inflation of *P* values around 0 emerges visibly when resampling at 50% of the full sample size.

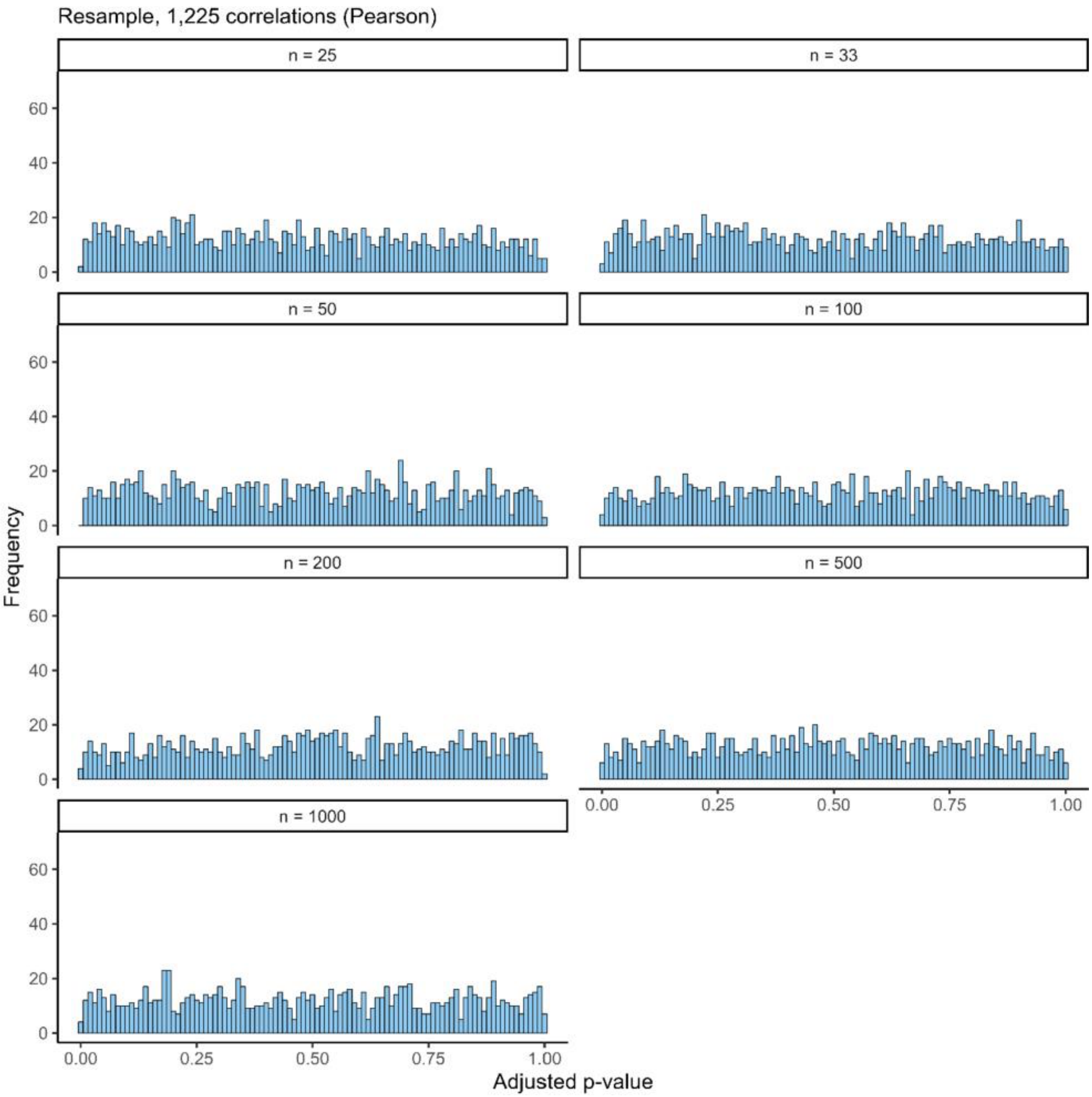


Supplementary Figure 2: **Distributions of adjusted *P* values under different resample sizes.** Note that *P* values are uniformly distributed across all resample sizes, as desired.

We show that we can correct inflation by adjusting P values, but unfortunately this does not lead to a reliable control of biased statistical power estimates. In subject level simulations with true effects (Supplementary Figure 3), if the discovery sample has low power the bias is under-corrected (Supplementary Figure 4), because random effects in the tails of the discovery are still more likely to be in the tails of a resample. On the other hand, if the discovery sample has high power the bias is over-corrected (Supplementary Figure 5), because true effects are less likely to be significant in a resample after adjusting *P* values.

**Subject level estimates of bias in statistical power under true effects**

In the main text we simulate null data at the subject level, with a brain measure and behavioural factor for each subject which is later correlated. Here we extend this approach to true effects by simulating non-gaussian multivariate data, so that simulated brain measures correlate with simulated behavioural factors across subjects. As this is relatively computationally expensive, in the main paper we take a computationally more efficient approach and simulate true data at the level of brain-behaviour correlations to consider a wider range of true effect scenarios.

To simulate data at the subject level, we generate two null samples as before (n = 1,000 subjects, resulting in 1,225 brain-behaviour correlations; see main text) but now further simulate 12 brain-behaviour correlations as true effects (in one case $\rho = 0.2$ and another case $\rho = 0.4$). The number of true effects here is arbitrarily chosen to be around 1% of all effects. To preserve the non-normal marginal distributions of brain connectivity estimates (edges being Pearson correlations bounded between -1 and 1) we simulate true effects via copula sampling (implementing methods described by Mathworks[2] in R). The main principle of copula sampling is to use a cumulative distribution function as a map between two given distributions, as this map conserves dependency structures[3]. We can then simulate data from a multivariate normal distribution (using the MASS package in R[4]) with a specified covariance structure (fixing $\rho = 0.2$ or $\rho = 0.4$ across all 12 true effects), lastly transforming the marginal distributions via the originally generated empirical null samples. We visualise the resulting data in Supplementary Figure 3.

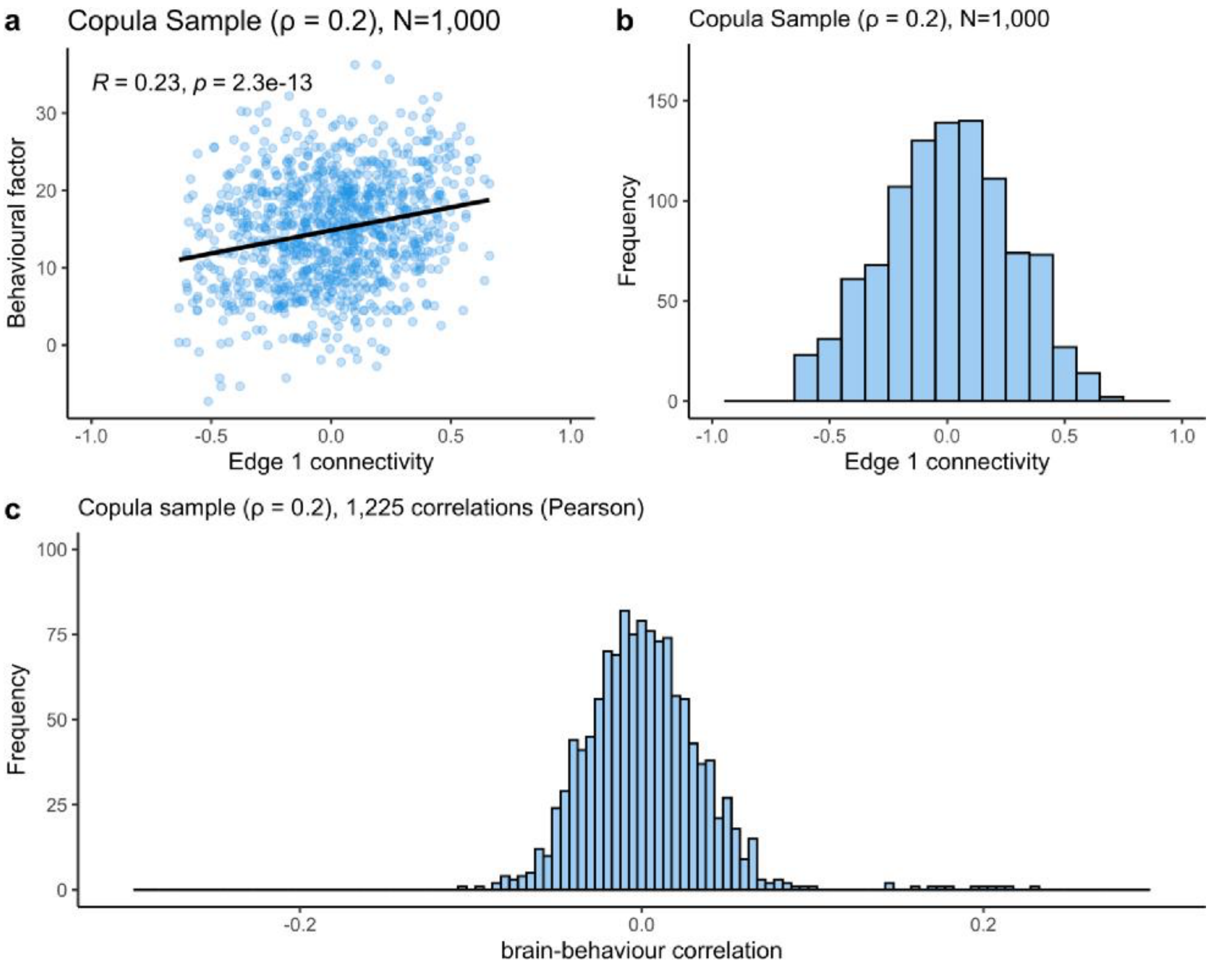


Supplementary Figure 3: **Visualising true effect simulations ($\rho$ = 0.2). a,** Scatterplot of behavioural factor against edge 1 connectivity across all participants ($n$ = 1,000). The associated Pearson correlation estimate is R = 0.23, $P < .001$. The black line indicates the simple linear regression line. **b,** We check that the copula preserves the marginal distribution of edge 1 across participants as non-gaussian (bounded between -1 and 1). **c,** Distribution of all 1,225 brain-behaviour correlations: most (1,213) of these are random, varying around 0, while the 12 simulated true effects vary around 0.2.

Using these true effect samples (simulated data containing 12 true effects at $\rho$ = 0.2 or $\rho$ = 0.4 respectively), we estimated statistical power under each significance threshold ($\alpha$ = .05, ... , $10^{-7}$) by the proportion of significant effects in the discovery sample which are significant again in a replication sample, averaged across iterations. We first obtained ground truth estimates by repeatedly generating true effect samples as replication samples (n = 100 iterations at each sample size). We then compared these results to estimations obtained when resampling from a discovery sample, as in Marek et al.[1]. We also generated a true effect sample with $n$ = 10,000 subjects and subsample up to 10% of full sample size ($n$ = 1,000). Lastly, we computed estimates obtained when resampling and adjusting $P$ values. Results for underlying true effects $\rho$ = 0.2 and $\rho$ = 0.4 are presented in Supplementary Figures 4 and 5 respectively.

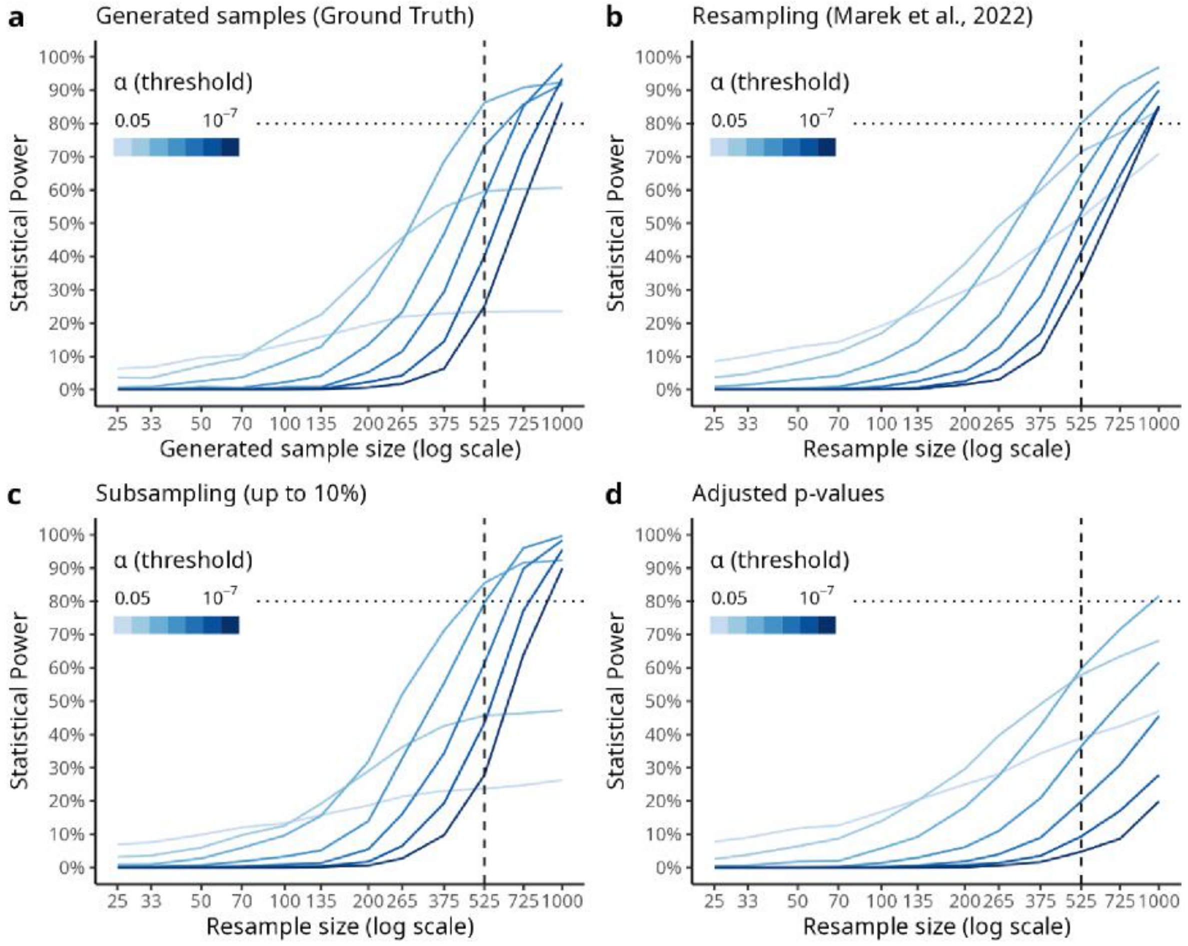


Supplementary Figure 4: **Statistical power estimates under true effects ($\rho$ = 0.2) a.** Ground truth of statistical power across sample sizes when 12 out of 1,225 brain-behaviour correlations with a true effect size of $\rho$ = 0.2. Notice that lenient significance thresholds ($\alpha$ = .05, .01) do not reach 80% power even with $n$ = 1,000 subjects, but corrected thresholds do ($\alpha$ = .001, ..., $10^{-7}$). This is because only 12 true effects were simulated, so at $\alpha$ = .05 around 61 (5% of 1213) random effects will be significant in the discovery sample, so the true (reproducible) effects will only make up a small proportion of significant effects in a replication sample under lenient thresholds. **b.** When estimating statistical errors using Marek et al.'s[1] methods we note that Bonferroni corrected estimates seem reliable, however uncorrected estimates are inflated (compared to **a**). **c.** When subsampling up to 10% of a large true effect sample ($n$ = 10,000), we note that estimates are reliable after Bonferroni correction, but also only slightly inflated for uncorrected estimates, providing the most reliable estimates. **d.** When resampling from a true sample ($n$ = 1,000) and computing adjusted $P$ values, we find that uncorrected error estimates are inflated while corrected error estimates are conservative, providing the least reliable statistical power estimates for true effects of size $\rho = 0.2$.

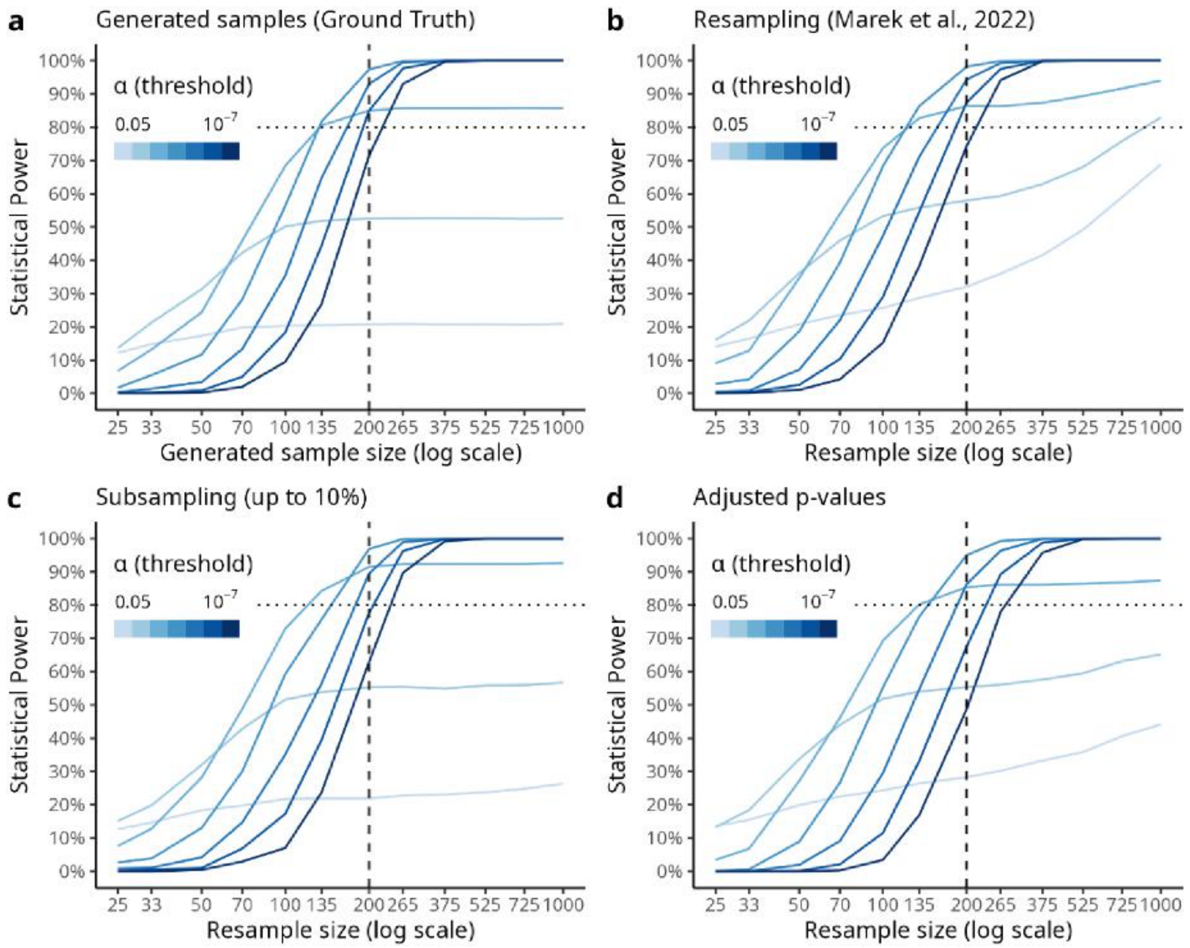


Supplementary Figure 5: **Statistical power estimates under moderate true effects ($\rho$ = 0.4). a,** Ground truth statistical power obtained when generating true effect samples ($\rho = 0.4$). Notice that lenient thresholds achieve lower power due to the small proportion of true effects (n = 12) compared to other effects which randomly pass the lenient thresholds. **b,** statistical power estimates obtained when resampling from a single true effect sample, as in Marek et al.[1]. Notice that statistical power is inflated for uncorrected estimates. **c,** Subsampling from a large true effect sample (n = 10,000) up to 10% of full sample size. We note that these estimates deviate little from the ground truth results (panel **a)**. **d,** Adjusting *P* values after resampling from a single true effect sample. We notice that for lenient significance thresholds, there is still some inflation of statistical power, whereas Bonferroni corrected estimates are conservative relative to ground truth (i.e., suggests larger sample sizes for similar power – the power curves are shifted to the right in panel d relative to panel a). For instance, to achieve 90% power, the ground truth results suggest using around 265 participants at alpha = $10^{-7}$, whereas with adjusted *P* values the estimate is closer to 375 participants.

Supplementary Figures 4 and 5 suggest that when underlying true effects ($\rho = 0.2$ and $\rho = 0.4$) are present in the sample, there is a noticeable difference between ground truth (generated) estimates and resampled estimates for uncorrected and lenient significance thresholds (compare panels **a** and **b** in Supplementary Figures 4 and 5). Subsampling up to 10% of full sample size gives reliable statistical power estimates when true effects are present (Supplementary Figures 4 and 5, panels **c**). On the other hand, we note that adjusting *P* values (Supplementary Figures 4 and 5, panels **d**) over-corrects Bonferroni estimates ($\alpha = .001, ..., 10^{-7}$), leading to conservative statistical power estimates relative to ground truth estimates.

While these subject level simulations respect the bounded distribution of brain connectivity estimates (non-gaussian distribution between -1 and +1), these simulations are still limited in how accurately they model real world data. One could consider alternative methods for simulating non-gaussian multivariate data[5,6] or more complete simulations of fMRI data[7,8]. However, we instead chose to simplify our simulations as done in the main text.
